# Simultaneous multi-transient linear-combination modeling of MRS data improves uncertainty estimation

**DOI:** 10.1101/2023.11.01.565164

**Authors:** Helge J. Zöllner, Christopher Davies-Jenkins, Dunja Simicic, Assaf Tal, Jeremias Sulam, Georg Oeltzschner

## Abstract

**Purpose:** The interest in applying and modeling dynamic MRS has recently grown. 2D modeling yields advantages for the precision of metabolite estimation in interrelated MRS data. However, it is unknown whether including all transients simultaneously in a 2D model without averaging (presuming a stable signal) performs similarly to 1D modeling of the averaged spectrum. Therefore, we systematically investigated the accuracy, precision, and uncertainty estimation of both described model approaches.

**Methods:** Monte Carlo simulations of synthetic MRS data were used to compare the accuracy and uncertainty estimation of simultaneous 2D multi-transient LCM with 1D-LCM of the average. 2,500 datasets per condition with different noise representations of a 64-transient MRS experiment at 6 signal-to-noise levels for two separate spin systems (scyllo-inositol and GABA) were analyzed. Additional datasets with different levels of noise correlation were also analyzed. Modeling accuracy was assessed by determining the relative bias of the estimated amplitudes against the ground truth, and modeling precision was determined by standard deviations and Cramér-Rao Lower Bounds (CRLB).

**Results:** Amplitude estimates for 1D- and 2D-LCM agreed well and showed similar level of bias compared to the ground truth. Estimated CRLBs agreed well between both models and with ground truth CRLBs. For correlated noise the estimated CRLBs increased with the correlation strength for the 1D-LCM but remained stable for the 2D-LCM.

**Conclusion:** Our results indicate that the model performance of 2D multi-transient LCM is similar to averaged 1D-LCM. This validation on a simplified scenario serves as necessary basis for further applications of 2D modeling.

## Introduction

Magnetic resonance spectroscopy (MRS) is the only non-invasive technique to investigate brain metabolism *in vivo*^1,2^. A typical conventional MRS measurement includes dozens to hundreds of transients that are subsequently averaged to produce a spectrum with sufficient SNR^3^. This spectrum is then approximated with a signal model, e.g., using 1D linear-combination modeling (LCM), to extract amplitude parameters.

An alternative to averaging and 1D modeling could be two-dimensional (2D) modeling of the entire dataset. 2D modeling methods were introduced over two decades ago^4–6^ to accommodate experiments like L-COSY^7^ and J-PRESS^8^ that encode information by deliberately changing conditions between transients. To retrieve this information, 2D modeling translates the relationship that connects the transients into explicit model coefficients. Dynamic MRS methods^9–14^ currently experience increased interest, for example, diffusion-weighted, functional, or multi-parametric MRS experiments. Furthermore, 2D modeling software is becoming increasingly accessible, e.g., through a dynamic fit module in FSL-MRS^10^ and the inclusion of the FitAID algorithm^6^ in the 2023 release of jMRUI (v7)^15^.

With renewed interest in and greater access to 2D modeling methods, it is important to validate and characterize their accuracy, precision, and uncertainty estimation, especially since even 1D modeling is strongly affected by decisions made during modeling, introducing substantial analytic variability^16–19^. Fortunately, today’s access to computing power and highly accelerated density matrix simulations^20–22^ allows for validation work that was not available when early 2D model algorithms were developed. This ability to precisely control the characteristics and quality of synthetic MRS data should be leveraged to validate emerging 2D methods.

In this study, we investigated whether 2D modeling of a conventional (1D) MRS experiment improves accuracy, precision, or uncertainty estimation compared to conventional (1D) modeling. To this end, we conducted Monte Carlo (MC) simulations^23,24^ of ideal synthetic MRS data and compared the results of simultaneous 2D multi-transient LCM to conventional 1D LCM of the averaged spectrum. 2,500 noise representations of a 64-transient MRS experiment with 6 signal-to-noise levels for two spin systems (scyllo-inositol and GABA) were modeled. Modeling accuracy was assessed by determining the relative bias, modeling uncertainty measured as standard deviations and Cramér-Rao Lower Bounds (CRLB).

## Methods

### Synthetic MRS data generator

We designed a synthetic MRS data generator based on the signal model used in the original Osprey LCM algorithm^25^. The generator accepts a basis set as input and applies separate amplitude, Lorentzian linebroadening, and frequency shift parameters for each basis function. Zero- and first-order phase, Gaussian linebroadening and an arbitrary convolution kernel are applied globally, i.e., to every basis function. Lastly, a baseline signal defined by a set of baseline amplitude parameters and parameterized as cubic splines or Voigt lines is added. Each parameter can be set up to a fixed value (including zero to ignore the parameter altogether) or drawn from a normal distribution with pre-defined mean and SD. Similarly, the set of metabolites to be included can be set. The user-supplied target amplitude-to-noise ratio for any peak in the spectrum is achieved by adding the required amount of uncorrelated Gaussian noise in the time domain. Synthetic data can be generated as single spectrum, multi-transient dataset, or with a parametrized relationship across an indirect dimension to simulate, e.g., echo time or diffusion-weighting series with exponential decays. The data is exported in NifTI-MRS for maximal compatibility^26^. The generator is available in the latest Osprey version (https://github.com/schorschinho/osprey).

### Monte Carlo simulations

30,000 synthetic spectra were generated for the main Monte Carlo MC simulation analysis. Separate spectra were simulated for two six-proton spin systems (scyllo-inositol (sI) and GABA) based on a sLASER sequence (TE = 30 ms)^27^. Six SNR levels (196, 128, 48, 12, 6, 3) were simulated, where SNR = 196 is comparable to a typical 2-ppm tNAA SNR^16^ and SNR = 6 is minimum singlet SNR cutoff suggested in a recent consensus paper^1^. The target SNR was defined for the averaged spectrum. The necessary noise power to achieve the target SNR was calculated based on the scyllo-inositol singlet peak and subsequently used for the GABA spin system as well, following the consensus noise definition of one standard deviation^28^. The following other parameters were set based on the distribution mean values from a prior study^16^: amplitude (0.15), Gaussian line broadening (5.70 Hz), and Lorentzian line broadening (2.42 Hz). All other model parameters were set to zero. For each scenario (2 metabolites × 6 SNR levels = 12 separate scenarios) 2,500 spectra were created, sufficient for the CRLBs to converge towards the standard deviation^29^.

In addition, 45,000 synthetic spectra at SNR 196 were simulated with correlated noise at 9 different correlation levels (correlation coefficient values r between 0.1 and .9), where the noise was correlated along the transient dimension. The amplitude and lineshape parameters were the same as for the other spectra.

Generalizability to other metabolites was tested in an additional analysis. A total of 42,500 synthetic spectra at SNR 196 were simulated with uncorrelated noise for 17 commonly found brain metabolites (ascorbate, aspartate, creatine, GABA, glycerophosphocholine (GPC), glutathione (GSH), Gln, Glu, mI, Lac, NAA, NAAG, phosphocholine, phosphocreatine, phosphoethanolamine (PE), sI, and taurine). The total signal – defined as the product of the number of protons and the amplitude parameter of the synthetic MRS data generator – was calculated for sI (6 protons and amplitude of 0.15) and held constant for all other metabolites. Another set of 42,500 synthetic spectra of these metabolites was generated at the same SNR level but using correlated noise with a correlation coefficient value of (r = 0.5) instead.

### Generalized linear-combination modeling

A new 2D-capable and highly customizable MATLAB-based LCM algorithm in Osprey (https://github.com/schorschinho/osprey/tree/MSM) was used for modeling. The least-squares optimization was performed on the real part in frequency domain (0.5 to 4 ppm) with the initial parameter guesses set to their respective ground truth values. The underlying models for data generation and modeling were matched; e.g., no baseline model was employed in the LCM to maximize the performance of CRLBs as an uncertainty estimator ^29^. For the 1D-LCM, the individual transients were averaged before modeling. For the 2D-LCM, all parameter estimates were fixed in the transient dimension, e.g., the same for all transients (**Figure 1**).

**Figure 1.**
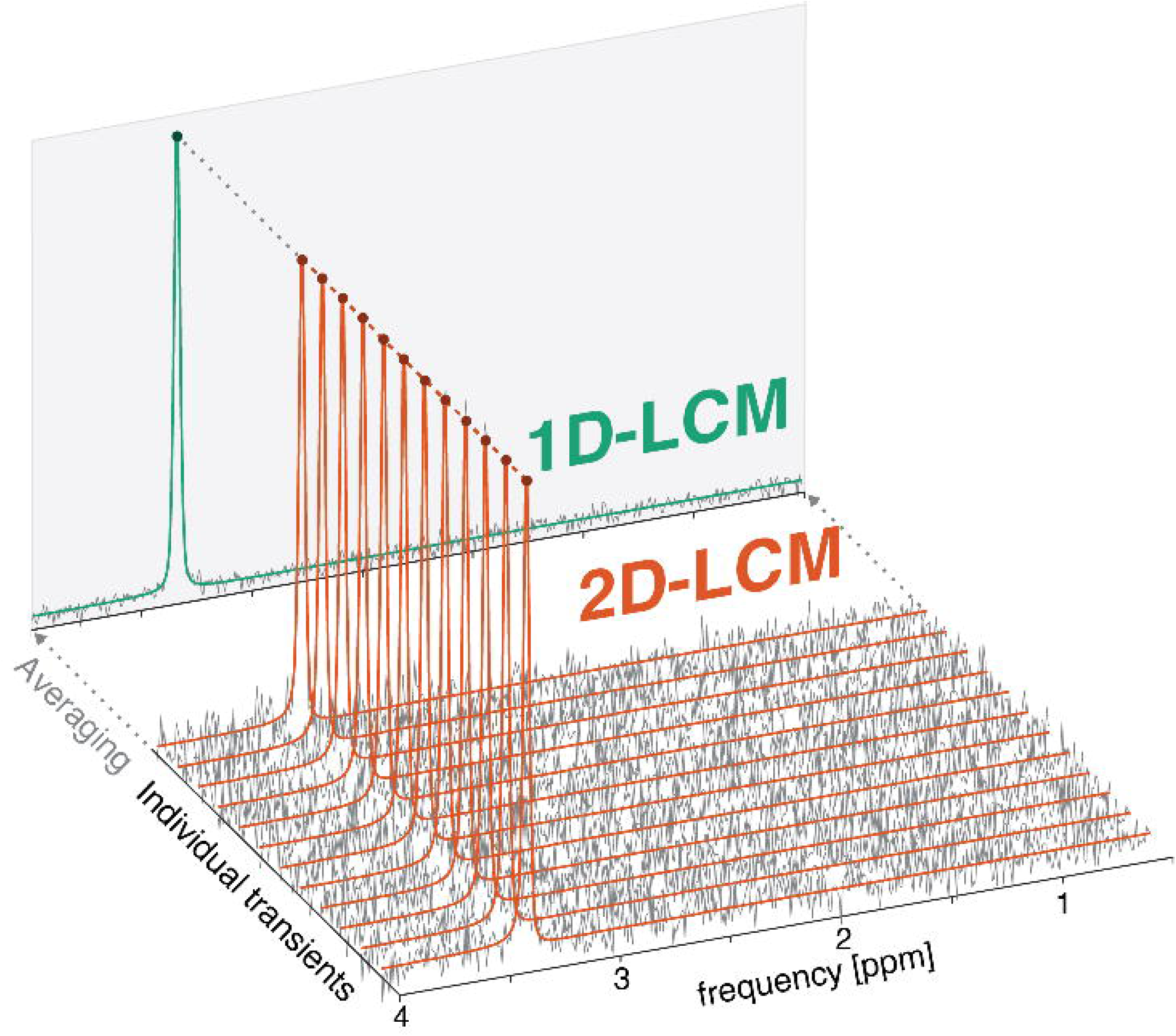
– Realization of 1D- and 2D-LCM on a 12-transient example dataset. For the 1D-LCM the individual transients are averaged (projected spectrum on the grey-shaded plane) prior to modeling (green). For the 2D-LCM, all individual transients are included in a single model (orange) and the model parameters are fixed along the transient dimension, e.g., the amplitude parameter is constrained to be the same for all transients.

### Model evaluation criteria

For each scenario the performance of the 1D- and 2D-LCM across the 2,500 synthetic spectra was evaluated as follows:

1. To assess model accuracy the relative amplitude estimation bias was calculated as the difference between the estimated and ground truth amplitudes, normalized by the ground truth amplitude (given in %). The standard deviation of the bias was used to assess model precision.
2. For the amplitude CRLBs the Fisher information matrix is formulated for each dataset. Under the assumption of independent and identically distributed random noise (Gaussian with zero mean) the (m,n)-th element of the Fisher matrix *F* for a model with n= 1,…N transients η= (η_1_(e),η_2_(e),…,η (e)) and m= 1,… model parameters e= (e_1_,e_2_,…e) is given by:

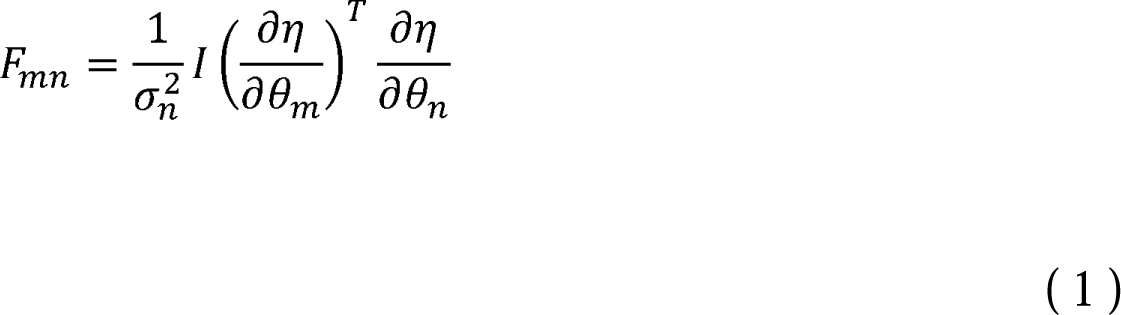

with a^2^ being the noise variance of the n-th transient. , being the Nx N identity matrix, and the partial derivatives with respect to the parameters (the Jacobian). The relative CRLBs are calculated based on the square-root of the trace of the inverted Fisher information matrix (using the Moore-Penrose pseudoinverse) and normalized by the amplitude estimate of the 1D- or 2D-LCM, respectively:

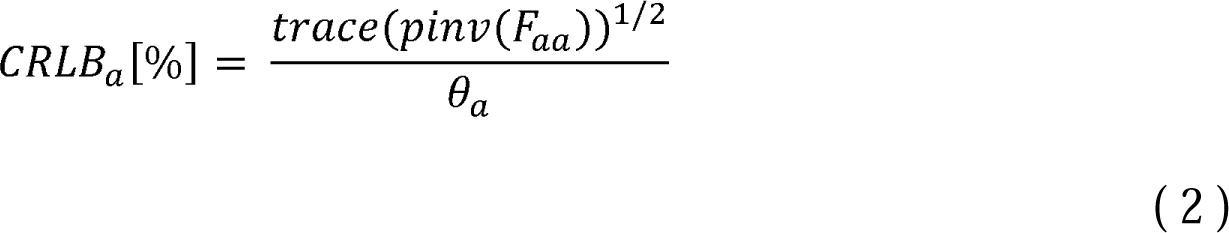

with index a referring to amplitude parameters e_a_. For the noise estimate a^2^, the signal-free frequency domain data between 8.5 and 12 ppm is first detrended using a linear fit. Afterwards the covariance matrix is calculated for the corrected real-valued data and the square-root of the trace is used as noise estimate a_n_. For the 1D-LCM (*n* = 1) this results in a single estimate. For the 2D-LCM, this results in a vector including one noise estimate per transient (*n* = number of transients). The CRLBs were used as an estimator of model uncertainty.
3. The true CRLBs were calculated by populating the Fisher information matrix with the ground-truth parameters e and the ground-truth noise amplitude.
4. Mixed CRLB estimates were calculated by either combining the ground-truth parameters, e, with the noise estimate, a_n_, or the ground-truth noise amplitude with estimated parameters, e. Thus, the impact of noise and parameter estimation on the CRLB can be investigated separately.
5. The standard deviation of the 2500 amplitude estimates of each scenario was calculated to validate the CRLBs. If the CRLBs are an estimator of the standard deviation, the ratio between mean CRLB and standard deviation should approach 1^29^. The CRLB-to-SD ratio was calculated for the estimated and the true CRLBs (see below). For the estimated CRLBs, cases with an estimated amplitude of zero (i.e., infinite CRLB) were removed from the SD and the mean CRLB calculations.

### Statistics

Differences in variance of the CRLBs between 1D- and 2D-LCM were tested with Fligner-Killeen’s test. Similarly, we calculated Pearson correlation coefficients to evaluate the agreement between 1D- and 2D-LCM for the amplitude estimates and CRLBs.

### Visualization

We used raincloud plots^30^ to visualize the amplitude bias where the boxplots included median, interquartile range (IQR), and whiskers (25^th^ or 75^th^ percentile ± 1.5 x IQR). Perfect reproduction of the ground truth is indicated by a dotted line. For the correlation analysis between the 1D- and 2D-LCM for the amplitude estimates the ground-truth amplitude is indicated by a straight line.

Two dotted lines mark a ±5% bias interval for the three highest SNR levels, and a ±25% bias interval for the lower SNR levels. Similarly, for the CRLB correlation analysis the true CRLB is indicated with a dotted line.

All code to reproduce the analysis can be found online^31^.

## Results

Visual inspection of randomly selected datasets indicated high performance for both the 1D- and 2D-LCM even for the low SNR scenario (see **Figure S1** in the Supporting Information).

### Amplitude estimation

The amplitude estimates for both model approaches are summarized in **Figure 2**. Most notably there is no difference in the standard deviation of the amplitude estimation for either model approach. The mean bias for both model approaches is negligible for the three highest SNR levels (<0.06% for sI and <0.98% for GABA). For the SNR level 6 (consensus-recommended cutoff value for a singlet SNR) a 4.6% ± 13.3% bias is found for sI and 15.2% ± 38.2% bias for GABA. Below this cutoff (SNR 3) the bias increases further to 7% ± 25% for sI and 47% ± 79% for GABA. The mean and standard deviation of the bias are generally higher for GABA than sI. It indicates that even for *in-vivo* realistic noise amplitudes the effect of signal height dominates the model performance even if the total integral of the spin systems is the same. The agreement between the amplitude estimates of both model approaches is almost perfect.

**Figure 2.**
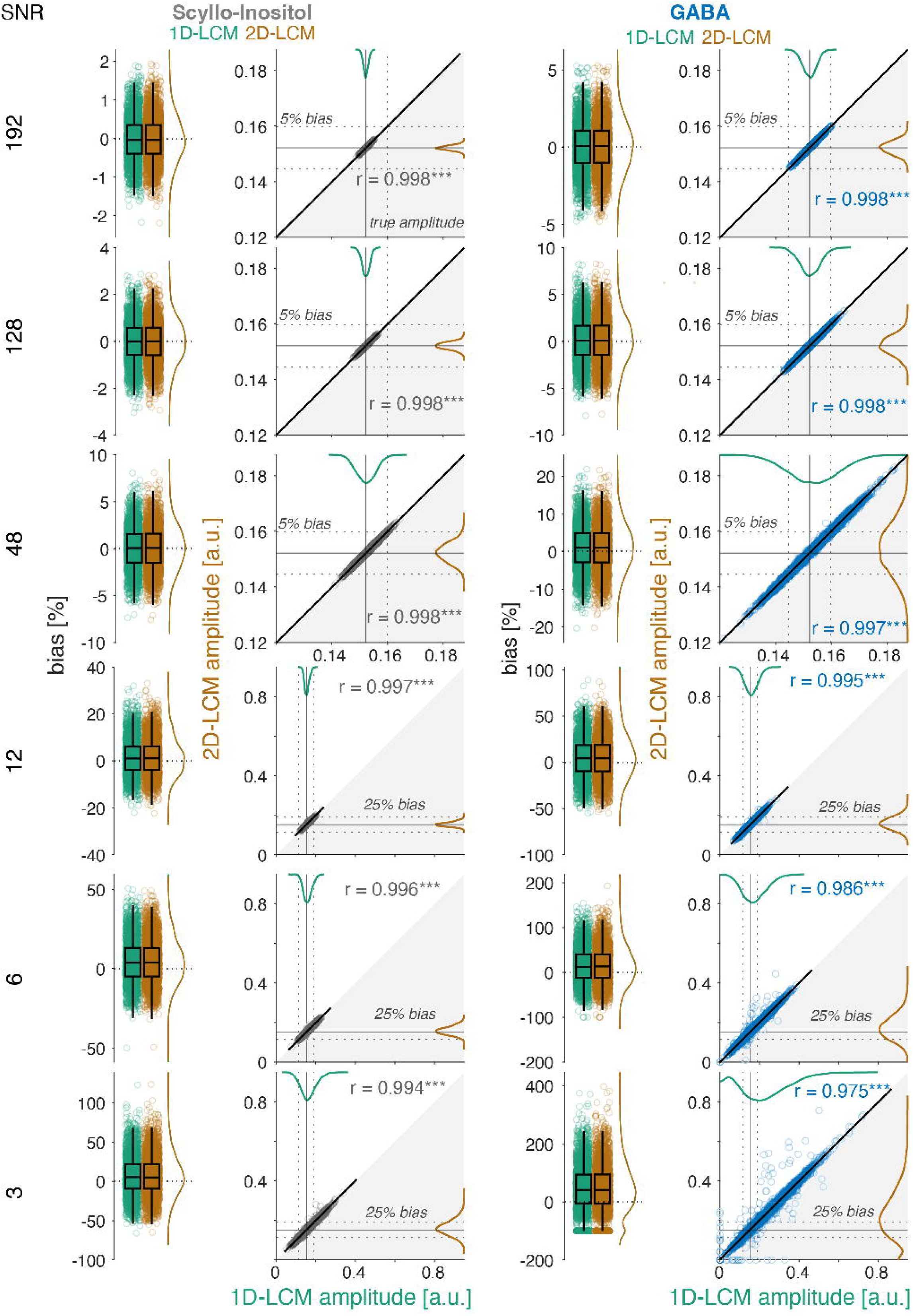
– Relative amplitude bias and amplitude estimate correlations of scyllo-inositol (left column) and GABA (right column) for 1D- (green) and 2D-LCM (orange) across six SNR levels (rows). Both models perform similarly in terms of mean and standard deviation of the relative bias and agree very well across all SNR levels. The relative amplitude bias is shown in the raincloud plots accompanied by the correlations between the amplitude estimates in the adjacent scatter plot. In the raincloud plots, the zero bias (perfect reproduction of the ground truth) is indicated by a dashed horizontal line. The correlation plots show individual datapoints and smoothed distributions; the ground-truth amplitude is indicated by straight horizontal and vertical lines. Two dotted lines mark a ±5% bias interval for the higher SNR levels (192, 128, 48) and a 25% bias interval for the lower SNR levels (12, 6, 3). The bisector is indicated by the gray shaded area.

### CRLB to standard deviation ratios

Ratios of all CRLB definitions (estimated noise SD / estimated parameters, true noise / true parameters, estimated noise SD / true parameters, true noise / estimated parameters) to the standard deviation across the 2500 amplitude estimates for all SNR levels are shown in **Figure 3**. Again, the closer this ratio is to 1, the better CRLBs perform as estimators of amplitude uncertainty due to random noise^29^. 2D-LCM has consistently lower CRLB-to-SD ratios compared to 1D-LCM, indicating a better performance as uncertainty estimator, although the effect is very small. For both metabolites, the CRLB-to-SD ratio is within 13% of unity for the three highest SNR levels, confirming that CRLBs are a good estimator of SD if the model perfectly describes the data and sufficient SNR is given. CRLB-to-SD ratios increase with decreasing SNR and the agreement between the true CRLB and the estimated CRLB decreases with decreasing SNR, confirming previous results^29^. The decrease in the CRLB-to-SD ratio for the 2D-LCM GABA SNR 3 scenario is due to the removal of zero-amplitude estimates. The fact that the true and estimated CRLB are always larger than the amplitude SD has previously been previously interpreted as an indicator that the estimated SD (from the noise) is a poor approximation for the true amplitude SD^29^.

**Figure 3.**
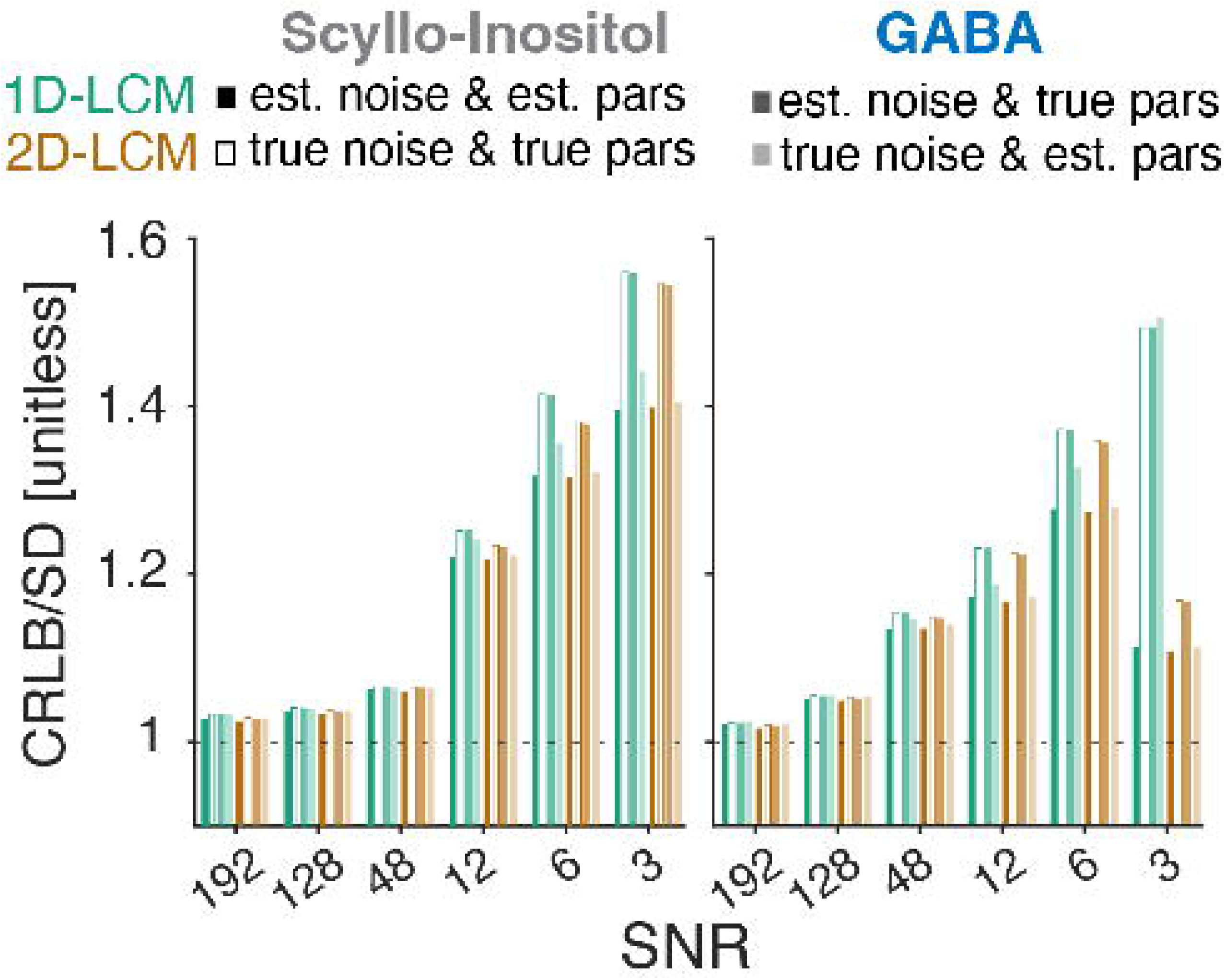
– CRLB to amplitude standard deviation ratio of scyllo-inositol (left) and GABA (right) for 1D-(green) and 2D-LCM (orange) across all SNR levels and CRLB definitions. The unity ratio is depicted as dotted line.

However, the mixed CRLBs calculated from the estimated noise and ground-truth parameters agree very well with the true CRLBs. In contrast, the mixed CRLBs combining ground-truth noise and estimated parameters agree less well with the true CRLBs and more with the estimated CRLBs. This indicates that the discrepancy is driven by how well the model parameters are estimated, and the quality of the noise estimation to not matter as much.

### CRLB estimation

**Figure 4** shows the correlation analysis of the CRLBs estimation for both model approaches and spin systems. Mean CRLBs do not significantly differ between 1D and 2D modeling and agree well with the true CRLBs. Interestingly, the standard deviation of the estimated CRLB for sI is significantly smaller for the 2D-LCM for the four highest SNR levels compared to the 1D-LCM. For GABA the 2D-LCM CRLB standard deviation is significantly smaller for the three highest SNR levels compared to the 1D-LCM. This is also reflected in the smaller correlation coefficient between the CRLBs of both models. The observed reduction in the standard deviation of the CRLBs is small, but means that 2D-LCM estimates the true uncertainty arising from random noise with slightly greater precision than 1D-LCM. The magnitude of this advantage diminishes as SNR decreases, as indicated by the slope of the regression line approaching the main diagonal. Similar figures for the mixed CRLBs are shown in the supporting information (**Figure S2** for estimated noise SD with ground-truth parameters and **Figure S3** for estimated parameters with ground-truth noise). **Figure S2** shows that 2D-LCM CRLBs have significantly smaller variance than 1D-LCM CRLBS for all SNR levels, indicating theoretically higher precision of the 2D-LCM uncertainty estimation. In contrast, **Figure S3** shows that 2D-LCM CRLBs have a significantly higher variance for perfect noise estimation than 1D-LCM CRLBs for all SNR levels except for sI at SNR = 3, indicating higher precision of the 1D-LCM uncertainty estimation if a high-quality noise estimate is included, for example, from a noise pre-scan.

**Figure 4.**
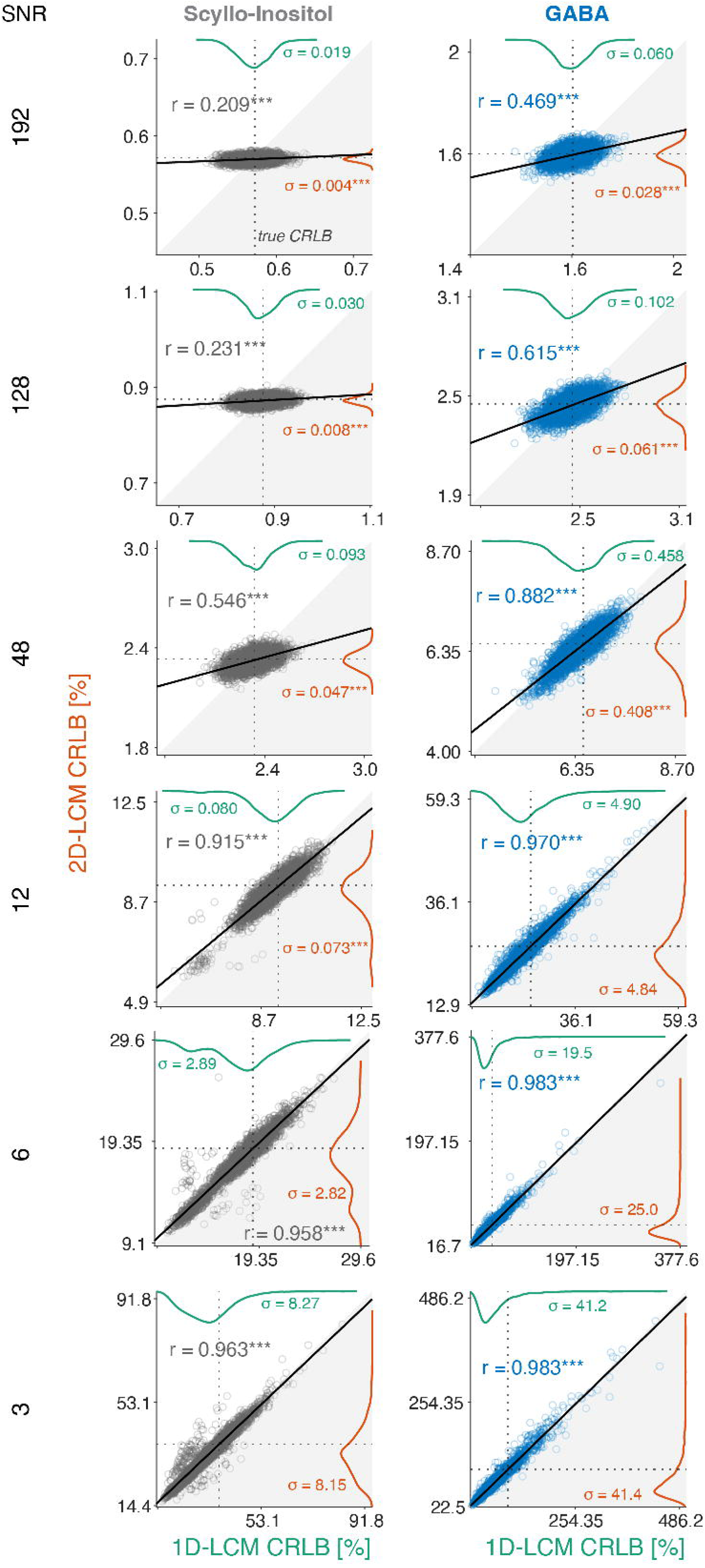
– CRLB estimates of scyllo-inositol (left column) and GABA (right column) for 1D-(green) and 2D-LCM (orange) across all SNR levels (rows). Mean CRLBs do not significantly differ between 1D- and 2D-LCM. However, the standard deviation of the 2D-LCM CRLBs is significantly smaller for SNR 192, 128, 48, and 12 for sI and 192, 128, and 48 for GABA compared to the 1D-LCM. This indicates that 2D-LCM estimates the true uncertainty arising from random noise with slightly greater precision than 1D-LCM. Dotted lines indicate true CRLB. The standard deviations CJ of the CRLB estimates are also reported (*** ≡ p < 0.001 for the Fligner-Killeen’s test).

### CRLB estimation for correlated noise

**Figure 5** shows the relative CRLB for both model approaches and spin systems for correlated noise with different correlation strengths. The mean relative CRLB for the 2D LCM is stable for all correlation levels r and the standard deviation is consistently small. For the 1D LCM, the mean and standard deviation of the relative CRLB decrease with decreasing correlation coefficients r. This is in line with earlier works showing improvements in CRLB estimates for correlated noise from different receiver coils^32^. To rule out any effects on the maximum-likelihood estimator during the optimization, we performed whiteness tests on a representative 1D noise vector of correlated (r = 0.5) and uncorrelated noise and found that both noise vectors represented additive white Gaussian noise.

**Figure 5.**
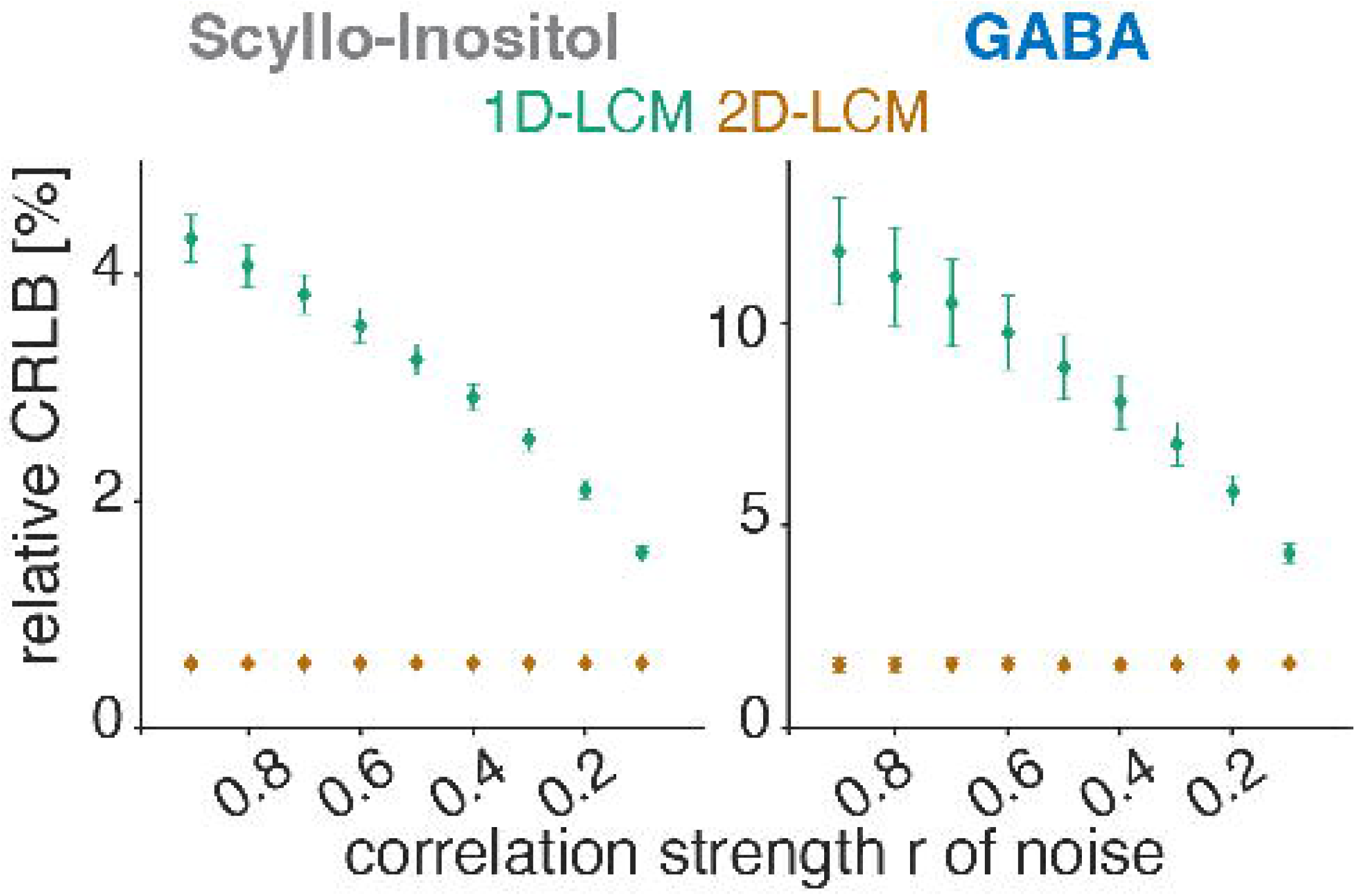
– CRLB estimates of scyllo-inositol (left column) and GABA (right column) for 1D-(green) and 2D-LCM (orange) for SNR 196 of correlated noise with different correlation coefficients r.

### Generalization to other metabolites

**Figure S4** summarizes the results across 17 metabolites. The mean and standard deviation for the amplitude bias agree well for both model approaches for all metabolites (**Figure S4 A**) and the agreement between the amplitude estimates is near-perfect (**Figure S4 B**). The ratio of the mean estimated CRLBs and true CRLB to the amplitude standard deviation reproduces well across metabolites and approaches 1 in all cases. Similarly, true CRLBs-to-SD ratios are always higher than the estimated CRLBs-to-SD ratios (**Figure S4 C**). The correlation between the CRLB estimates of both model approaches ranges between 0.2 and 0.6 for different metabolites (**Figure S4 D**). Weaker correlations coincide with lower standard deviations of CRLB estimates of the 2D-LCM. Metabolites with strong singlet signals exhibit lower correlation and smaller standard deviations of the CRLB estimates of the 2D-LCM, while this advantage diminishes for highly J-coupled metabolites. The relative CRLB is generally smaller for the 2D-LCM in case of correlated noise (**Figure S4 E**). Overall, the results generalize well across different metabolites.

## Discussion

This study aimed to investigate whether 2D multi-transient LCM of conventional 1D MRS data improves accuracy, precision, and uncertainty estimation compared to conventional 1D-LCM of averaged spectra. We deliberately designed an idealized, well-controlled representation of a multi-transient MRS experiment to allow the most precise characterization of the 2D-LCM algorithm. This work can serve as a benchmark scenario and can be adapted to validate more complex scenarios, for example, by adding frequency- and phase drift, as well as *in-vivo* realistic metabolite compositions and their dynamic changes.

We found that 1D- and 2D-LCM achieved comparable accuracy and precision of amplitude estimation, with no significant advantage offered by either. Uncertainty estimation was delivered with greater precision by 2D multi-transient LCM, evidenced by the substantially reduced (up to 79%) standard deviation of the estimated CRLBs, without biasing them away from the true CRLBs. Therefore, 2D-LCM estimates the true uncertainty arising from random noise better. It should be noted that the practical benefit was rather small.

Notably, both methods exhibited a positive amplitude estimation bias, i.e., on average, the estimated amplitude was higher than the ground truth. This bias increased in low-SNR scenarios, and likely arises from the metabolite amplitude parameter constrained to be non-negative, which is commonly used in linear-combination modeling algorithm, and the fact that only positive amplitudes were included in the simulation.

It is possible that improved model precision in 2D-LCM is only achieved when information is different between the transients. This was the case in the recent study of 2D modeling of synthetic, overlapping peaks in a multi-TE series^11^; in this case, precision improves due to the reduced model freedom obtained through re-parametrization and the concurrent constraints on parameter estimation. In contrast, we studied an idealized 1D-multi-transient experiment where the signal amplitude was not modulated between transients. While we anticipated benefit from including all noise representations (instead of just the average noise), this advantage was marginal. This suggests equivalence of our 1D- and 2D least-squares optimization procedures and indicates that the noise variance term in the 1D-CRLB expressions sufficiently approximates the main diagonal noise variance estimates from the 2D-CRLB noise correlation matrix. The true benefits of noise estimation with 2D-LCM may only become apparent for scenarios in which noise is, in fact, correlated, e.g., when simultaneously modeling signals from different, potentially noise-correlated, receiver coils, as previously shown^32^ or when Brownian motion in the voxel is considered as the source of the noise^33^. The second set of synthetic data used in this study confirmed that 2D-LCM effectively separates correlated and uncorrelated noise components, leading to stable CRLB estimation for all tested correlation coefficient values r. In comparison, the mean CRLB from the 1D-LCM was substantially increased as the estimated noise from the averaged 1D spectrum does not allow for this separation.

For both models, we found CRLBs to be reliable uncertainty estimators of the standard deviation, across multiple identical measurements if SNR was reasonably high (in this study SNR 48), indicated by the CRLB-to-SD ratio approaching 1. This confirms previous work^29^ showing that CRLBs adequately quantify aleatoric model uncertainty, i.e., uncertainty due to random measurement noise, as long as the model accurately describes the data, which was certainly the case in this perfectly controlled ideal scenario. The CRLB-to-SD ratio for 2D-LCM was consistently (but only slightly) lower than for 1D-LCM, again suggesting that the inclusion of all transients benefits the noise estimation, but only achieving a small effect in practice.

New experimental paradigms in functional, diffusion-weighted and multi-parametric MRS have sparked a renewed interest in 2D modeling of MRS data^9–12^. While classic algorithms like ProFit^4,5^ never broke into mainstream use, the accelerated rate of innovation in MRS analysis software makes 2D modeling accessible to a wider audience, e.g., with FitAID ^6^ integrated into jMRUI and a 2D modeling framework in FSL-MRS^10,34^. This resurgence should be accompanied by thorough validation and benchmarking, as it has become clear that even 1D model procedures are very susceptible to small changes in algorithmic configuration^16–19^.

Validation can be achieved with precisely controlled synthetic data. Interest in and availability of synthetic data generation is further motivated by the need for realistic training data for MRS machine learning methods^35–38^. Synthetic MRS data generators are currently predominantly created for in-house use and not often made available as open-source tools. It will be important for the MRS field to reach consensus on what constitutes ‘realistic’ synthetic data, if reproducibility and validity of current and future analysis methods are to be evaluated fairly.

Discussion is ongoing, e.g., in a recently founded working group of the Code and Data Sharing Committee of the ISMRM MRS study group. The synthetic MRS data generator presented here may serve as a template as it accepts any metabolite basis set for input, draws from parameter distributions for all physical model parameters, can create interrelated MRS data, and generates the synthetic output in NIfTI-MRS format^26^ for maximal interoperability.

### Limitations and Future Directions

This study uses an idealized, simplified scenario to study theoretical improvements in model accuracy and uncertainty estimation of 2D modeling. While the findings generalize well across different metabolites, future studies need to evaluate the applicability to in-vivo data by progressively approaching *in-vivo-*like conditions. Real-life effects like an increasing number of overlapping signals at *in-vivo* realistic concentrations, frequency- or phase-drift^39^, amplitude fluctuations considered during fMRS^10–12^ or due to motion, and phase cycling need to be carefully designed to recreate sources of transient-to-transient variability. The ultimate application to *in-vivo* data where the ground truth for all physical parameters is unknown should be done with caution unless their impact is properly understood.

## Conclusion

2D multi-transient LCM of conventional 1D-MRS data is comparable to 1D-LCM of the averaged spectrum in terms of accuracy and precision, with small benefits of uncertainty estimation for uncorrelated noise and substantial benefits for correlated noise. The presented framework can be used for the validation of 2D modeling with well-controlled synthetic data for more complex applications of 2D modeling.

## Acknowledgement

This work has been supported by NIH grants R00 AG062230, R21 EB033516, K99 AG080084, R01 EB016089, R01 EB023963, and P41 EB031771.

## Data and code availability statements

The code to run the analysis is available on Center for Open Science at https://osf.io/k9hzx/

## Supporting Information Captions

*Figure S1 – Example of 1D- (green) and 2D-LCM (orange) of multi-transient MRS for different SNR levels for scyllo-inositol (gray) (A) and GABA (blue) (B). For easier visualization the residual was only included for the 1D-LCM of the averaged spectra.*

*Figure S2 – CRLBs using estimated noise SD and ground-truth parameters of scyllo-inositol (left column) and GABA (right column) for 1D- (green) and 2D-LCM (orange) across all SNR levels (rows). Mean CRLBs do not significantly differ between 1D- and 2D-LCM. However, the standard deviation of the 2D-LCM CRLBs is significantly smaller for all SNR levels compared to the 1D-LCM. This indicates that 2D-LCM estimates the true uncertainty arising from random noise with slightly greater precision than 1D-LCM. Dotted lines indicate true CRLB. The standard deviations* CJ *of the CRLB estimates are also reported (*** ≡ p < 0.001 for the Fligner-Killeen’s test)*.

*Figure S3 – CRLBs using ground-truth noise and estimated parameters of scyllo-inositol (left column) and GABA (right column) for 1D- (green) and 2D-LCM (orange) across all SNR levels (rows). Mean CRLBs do not significantly differ between 1D- and 2D-LCM. Dotted lines indicate true CRLB. The standard deviations CJ of the CRLB estimates are also reported (** ≡ p < 0.05; *** ≡ p < 0.001 for the Fligner-Killeen’s test).*

*Figure S4 – Generalizability analysis across 17 metabolites at SNR level 196 with uncorrelated noise (A-D) and correlated noise at correlation strength r=0.5. (A) Relative amplitude bias for 1D- (green) and 2D-LCM (orange). (B) Amplitude estimates correlation coefficient between 1D- and 2D-LCM. (C) Mean CRLB to amplitude standard deviation. (D) CRLB correlation coefficient between 1D- and 2D-LCM. (E) CRLB estimates for 1D- (green) and 2D-LCM (orange) for correlated noise at SNR level 196. The analysis indicates good generalizability of the findings across different metabolites.*

