## Supplementary Material for "Simultaneous multi-transient linear-combination modeling of MRS data improves uncertainty estimation"

### Supporting Information

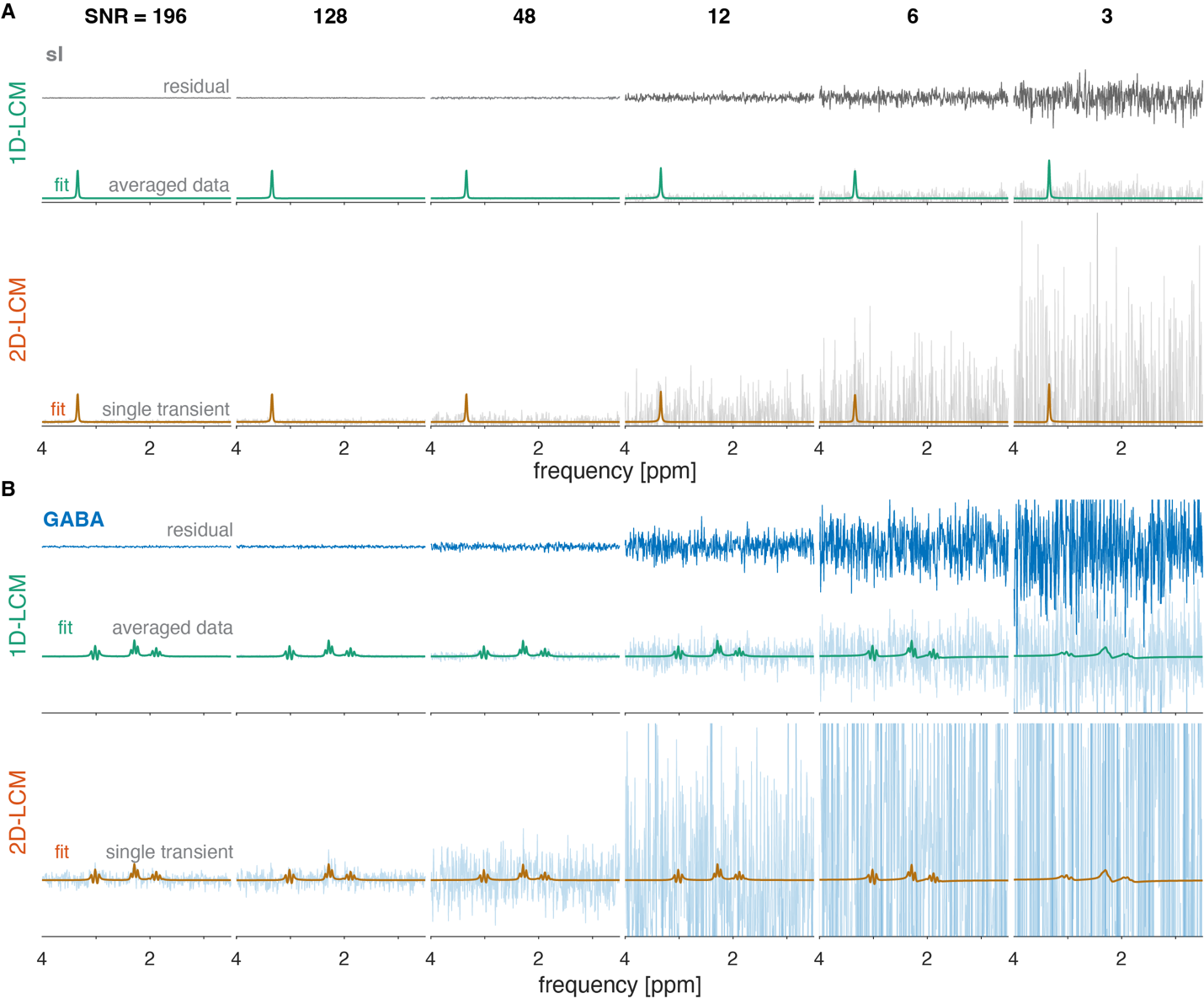

Figure S1 – Example of 1D- (green) and 2D-LCM (orange) of multi-transient MRS for different SNR levels for scyllo-inositol (gray) (A) and GABA (blue) (B). For easier visualization the residual was only included for the 1D-LCM of the averaged spectra.

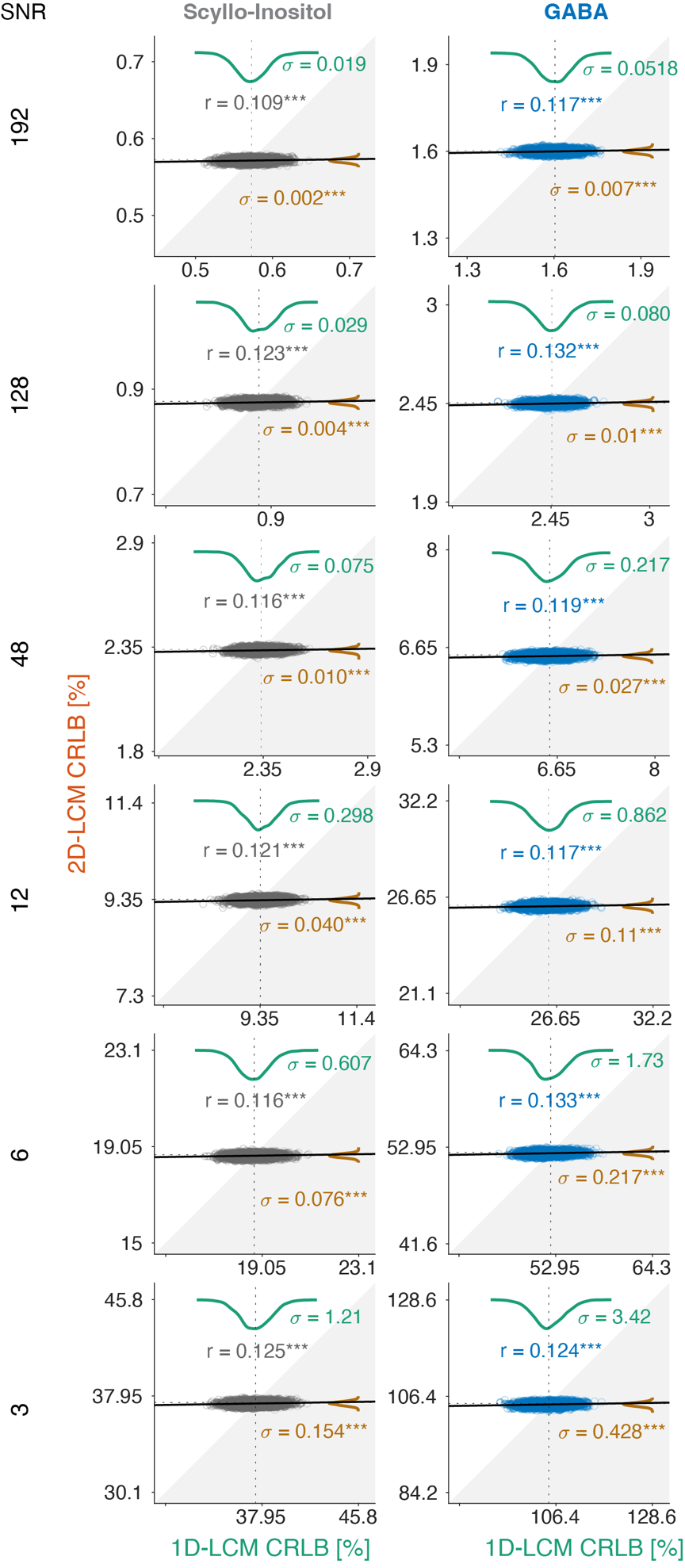

Figure S2 – CRLBs using estimated noise SD and ground-truth parameters of scyllo-inositol (left column) and GABA (right column) for 1D- (green) and 2D-LCM (orange) across all SNR levels (rows). Mean CRLBs do not significantly differ between 1D- and 2D-LCM. However, the standard deviation of the 2D-LCM CRLBs is significantly smaller for all SNR levels compared to the 1D-LCM. This indicates that 2D-LCM estimates the true uncertainty arising from random noise with slightly greater precision than 1D-LCM. Dotted lines indicate true CRLB. The standard deviations $\sigma$ of the CRLB estimates are also reported (*** ≡ p < 0.001 for the Fligner-Killeen’s test).

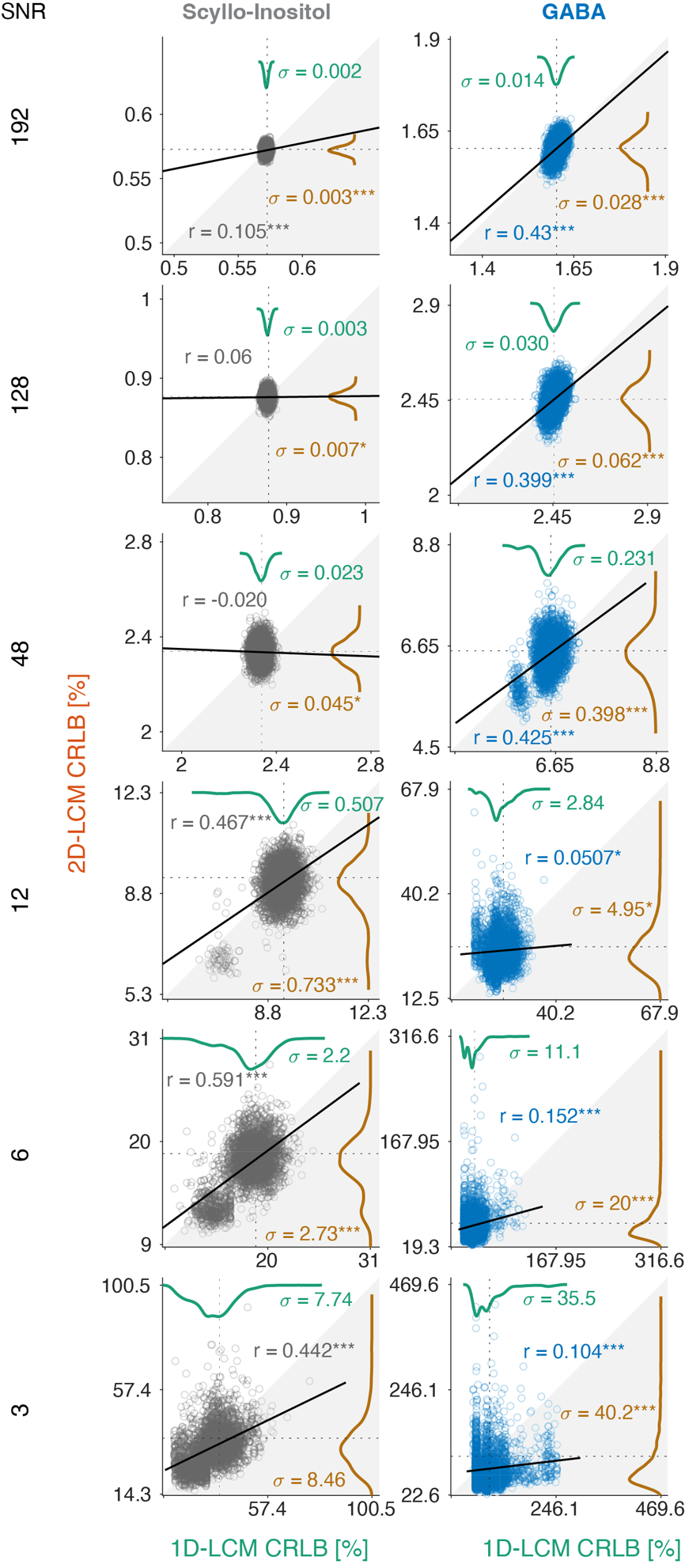

Figure S3 – CRLBs using ground-truth noise and estimated parameters of scyllo-inositol (left column) and GABA (right column) for 1D- (green) and 2D-LCM (orange) across all SNR levels (rows). Mean CRLBs do not significantly differ between 1D- and 2D-LCM. Dotted lines indicate true CRLB. The standard deviations $\sigma$ of the CRLB estimates are also reported (* ≡ p < 0.05; *** ≡ p < 0.001 for the Fligner-Killeen’s test).

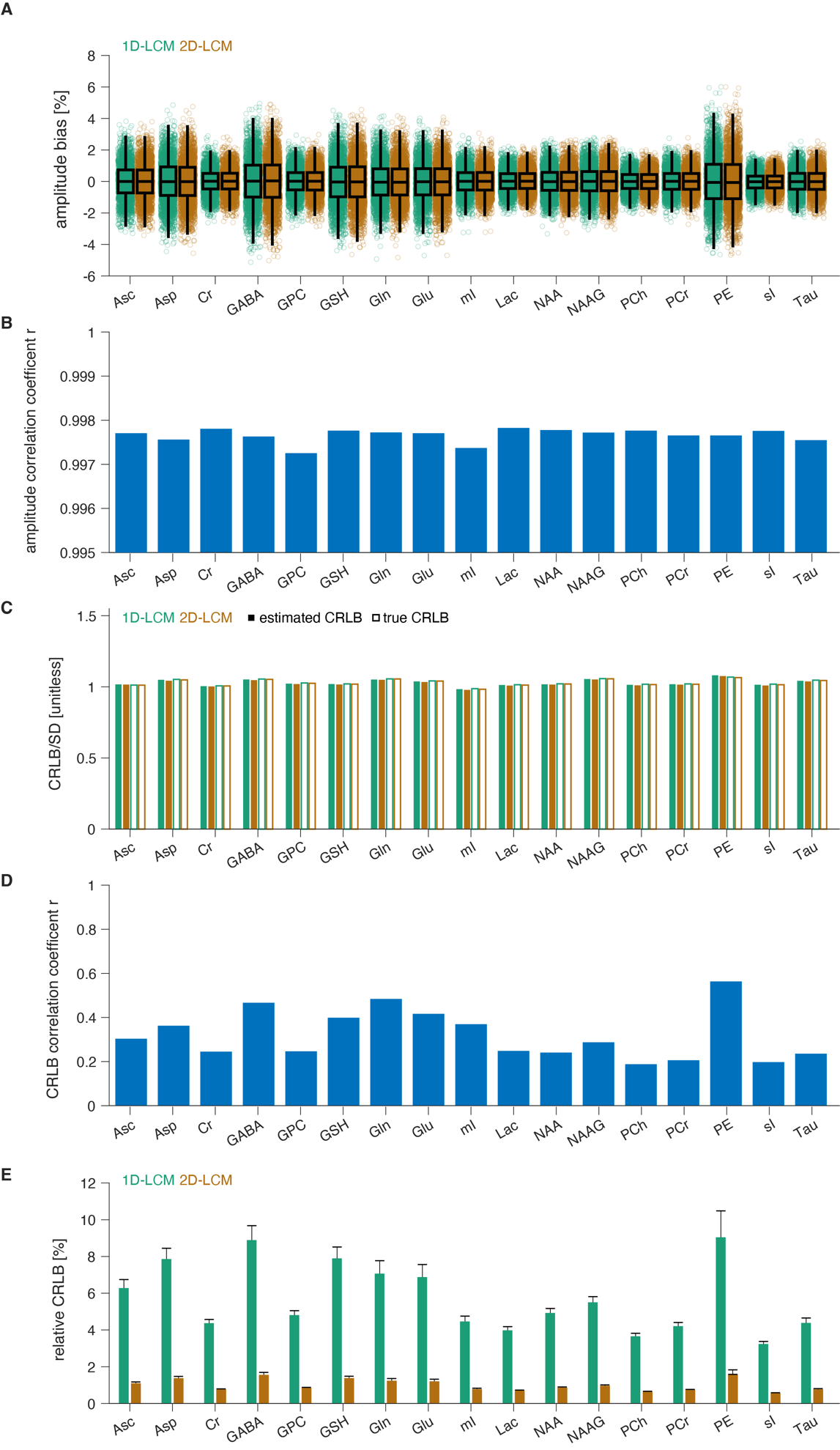

Figure S4 – Generalizability analysis across 17 metabolites at SNR level 196 with uncorrelated noise (A-D) and correlated noise at correlation strength r=0.5. (A) Relative amplitude bias for 1D- (green) and 2D-LCM (orange). (B) Amplitude estimates correlation coefficient between 1D- and 2D-LCM. (C) Mean CRLB to amplitude standard deviation. (D) CRLB correlation coefficient between 1D- and 2D-LCM. (E) CRLB estimates for 1D- (green) and 2D-LCM (orange) for correlated noise at SNR level 196. The analysis indicates good generalizability of the findings across different metabolites.
